# Pulse-Width Modulation-based TMS mimics effects of conventional TMS on human primary motor cortex

**DOI:** 10.1101/2021.11.24.469832

**Authors:** Majid Memarian Sorkhabi, Karen Wendt, Jacinta O’Shea, Timothy Denison

**Affiliations:** MRC Brain Network Dynamics Unit, Nuffield Department of Clinical Neurosciences, University of Oxford, Oxford, OX1 3TH, UK; Wellcome Centre for Integrative Neuroimaging (WIN), Oxford Centre for Human Brain Activity (OHBA), University of Oxford Department of Psychiatry, Warneford Hospital, Warneford Lane, Oxford, UK; Department of Engineering Science, University of Oxford, Oxford, OX1 3PJ, UK

**Author notes:** **Corresponding author:** Majid Memarian Sorkhabi, Postal address: Institute of Biomedical Engineering, Old Road Campus Research Building, Oxford, UK OX3 7DQ.

**Keywords:** Transcranial magnetic stimulation, programmable TMS, TMS pulse generator, MEP measurement

## Abstract

**Objective:** We developed a novel transcranial magnetic stimulation (TMS) device to generate flexible stimuli and patterns. The system synthesizes digital equivalents of analog waveforms, relying on the filtering properties of the nervous system. Here, we test the hypothesis that the novel pulses can mimic the effect of conventional pulses on the cortex.

**Approach:** A second-generation programmable TMS (pTMS2) stimulator with magnetic pulse shaping capabilities using pulse-width modulation (PWM) was tested. A computational and an in-human study on twelve healthy participants compared the neuronal effects of conventional and modulation-based stimuli.

**Main results:** Both the computational modeling and the in-human stimulation showed that the PWM-based system can synthesize pulses to effectively stimulate the human brain, equivalent to conventional stimulators. The comparison includes motor threshold, MEP latency and input-output curve measurements.

**Significance:** PWM stimuli can fundamentally imitate the effect of conventional magnetic stimuli while adding considerable flexibility to TMS systems, enabling the generation of highly configurable TMS protocols.

**Highlights:**

- The PWM method promises the implementation of flexible neurostimulation
- PWM magnetic pulses were well tolerated by the participants without adverse events
- RMTs and MEPs were compared for PWM and conventional stimuli
- PWM-equivalent of conventional pulses has relatively similar effects on the cortex
- The use of digital synthesis techniques to create novel patterns is a promising method for future neuromodulation

## Introduction

Transcranial magnetic stimulation (TMS) is a non-invasive method utilized to stimulate and modulate the nervous system. Most TMS devices are limited to predefined pulse shapes, only generating either monophasic or biphasic cosine-shaped pulses. Repetitive TMS protocols, particularly monophasic paradigms, have always been associated with an energy recovery challenge [1]. Recently, the use of state-of-the-art power electronic instruments has permitted more control over the waveform parameters [2] [3] [4]. A novel technique utilizing pulse width modulation (PWM), called programmable TMS or pTMS [5], enables the imitation of a wide range of arbitrary pulses. This structure can generate PWM-equivalents of monophasic, biphasic and polyphasic pulses with low interstimulus intervals (1 ms) by optimally recovering the energy delivered to the coil.

This study introduces a first-in-human study which uses the second generation of the pTMS device (pTMS2), making use of the modular device topology. To validate the effect of this device, the conventional monophasic pulse of a Magstim 200^2^ stimulator was imitated by the pTMS2 device. Computational modelling, resting motor thresholds (RMT), motor evoked potential (MEP) amplitude, latency and input-output (IO) curve measurements were compared for both devices.

## Materials and Methods

### TMS devices

We used a custom-built pTMS2 device that cascades two of the inverter cells introduced in [5] [6] [7] and generates magnetic pulses with five voltage levels. The PWM can approximate any reference waveform, but the pulse will include the fundamental harmonic of the reference pulse as well as its higher frequency harmonics. With this principle, pulses of different shapes and lengths can be generated as single pulses and trains of pulses (Fig S1-S4).

The conventional monophasic pulses were generated with a commercial Magstim 200^2^ (Magstim Co., UK). Both devices were connected to the same 70 mm figure-of-eight coil (Magstim Co., P/N 9925–00) with an adapter (Magstim Co., P/N 3110–00). The output pulses of the two devices are shown in Figure 1a.

**Figure 1.**
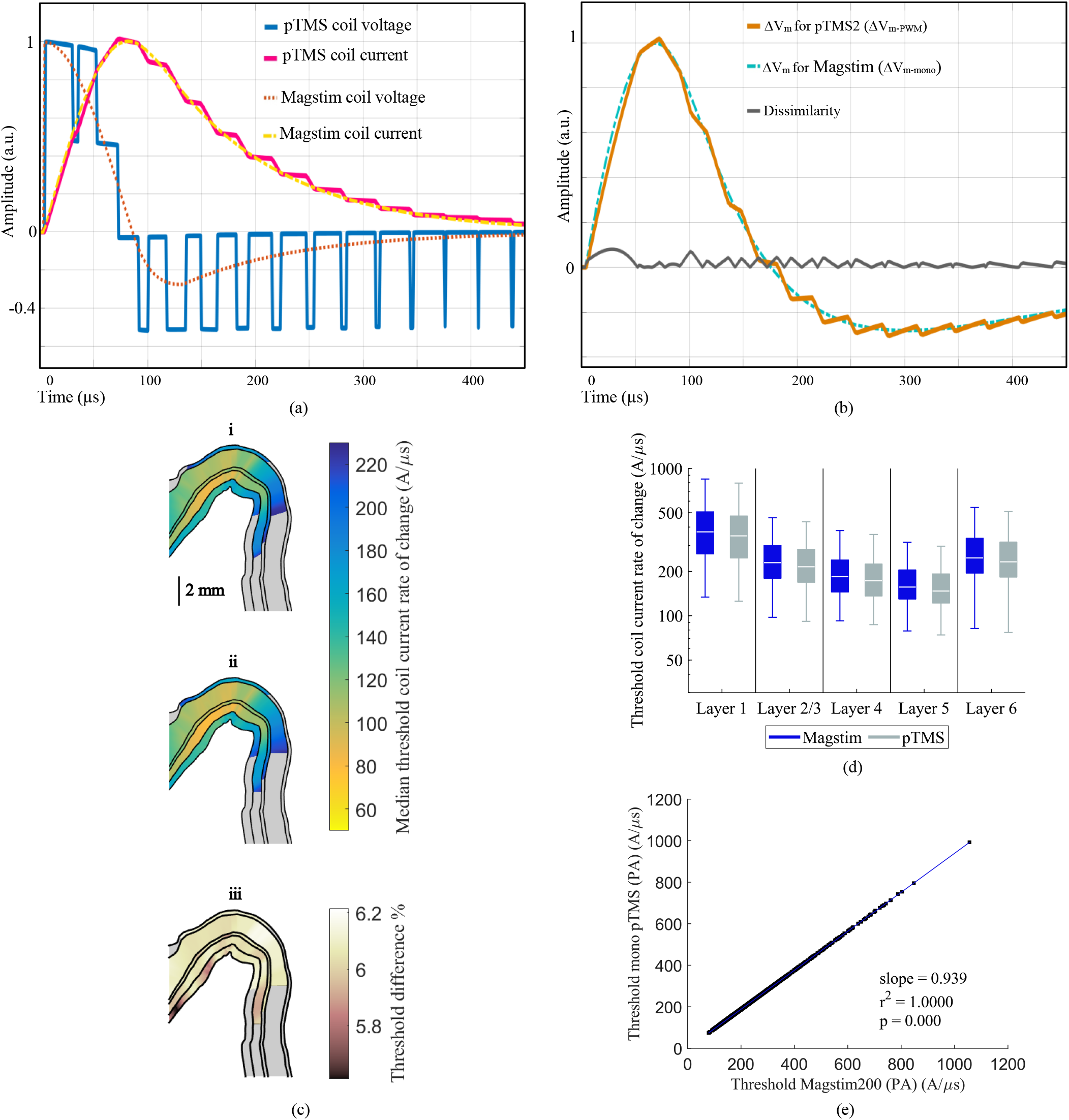
Comparison of the Magstim 200^2^ and pTMS2 system outputs and modelled responses for monophasic stimuli. (a) PWM coil voltage and current waveforms generated in the pTMS2 architecture in comparison with the Magstim waveforms. (b) Expected voltage changes in the membrane (ΔV_m_) from the RC model, when applying the Magstim 200^2^ and the pTMS2 stimuli. The dissimilarity of the two membrane voltage changes is defined as |Δ*V_m–PWM_* – Δ*V_m–mono_*|. 150 μs was selected as the nerve membrane time constant under magnetic stimulation to predict ΔVm in response to magnetic stimulation [22]. Based on this modeling, the difference between the devices would be expected to be less than 8%. (c) The median thresholds of the change in coil current for the six cortical layers shown on a 2D cross section of the crown of the pre-central gyrus for the pulse waveforms from (i) the Magstim 200^2^ stimulator and (ii) the pTMS2 device. (iii) shows the modelled percent difference in median thresholds between the Magstim and pTMS2 pulses. (d) The modelled neural activation thresholds within the cortical area representing the hand muscle are shown in log scale, where blue represents the data from the Magstim 200^2^ and grey from the pTMS2. Each boxplot includes the data from five clones with the outliers removed. (e) Correlation between the threshold coil current rate of change for the two pulses, with the linear regression displayed in blue.

## Physiological Response Models

To understand how the PWM stimuli, with their high-frequency harmonics, interact with neural tissues, two biophysically based models were applied before conducting the in-human study:

### RC model

Considering only the subthreshold dynamics of the neuronal membrane, a resistor–capacitor (RC) model can estimate the membrane potential variation (ΔV_m_), where ΔV_m_ biophysically outlines the shift of the membrane potential from the resting state of the membrane [8]. This model approximates the membrane as a low-pass filter with a time constant of 150 μs.

### Morphological neural models

A model which integrates morphological neural models with transcranially induced electric fields is used to compare the neural response to the Magstim and pTMS2 pulses [9] [10], similar to a previous study [11](see supplementary file for more details). The Simulink models for the temporal waveforms were adjusted to replicate the stimulation pulses of the devices used in the in-human study.

## In-human study

### Participants

Twelve healthy participants (mean age: 28.6 years, range: 22–37 years; 4 male) gave their informed consent to participate in the study which was approved by the Central University Research Ethics Committee (CUREC), University of Oxford (R75180/RE002). All participants were right-handed as assessed by the Edinburgh Handedness Inventory (Oldfield, 1971), had no current significant medical condition and reported no other contraindications to TMS.

### Procedure

Within each session, conventional and PWM stimuli were applied using the Magstim stimulator and pTMS2 devices, respectively, in counterbalanced order. The participants were seated in a chair with their arms resting on a pillow on top of a table in front of them. The coil was positioned over the left primary motor cortex and oriented at 45° to the midline with the handle pointing backwards. At the beginning of each session, the motor hotspot was determined with the Magstim stimulator, which indicates the optimal scalp position where MEPs could be elicited in the right first dorsal interosseous (FDI) muscle. ABrainsight neuronavigation system (Rogue Research Inc., Montreal, Canada) was used to track the position and orientation of the coil.

Electromyography (EMG) was recorded from the FDI of the right hand by positioning disposable neonatal ECG electrodes (Henley’s Medical, Welwyn Garden City, UK) in a belly-tendon montage, with the ground electrode over the ulnar styloid process. The RMT, defined as the minimum intensity required to evoke MEPs with ≥50 μV peak-to-peak at rest in 5 out of 10 trials [12], was measured and compared for both devices.

For the IO curve, MEPs at intensities up to the maximum voltage achievable by the pTMS2 device (see limitations section) were measured. Similar to other recent studies [13], TMS stimuli were applied in increasing order from low to high intensities in steps of 3% of the maximum stimulator output (MSO) of the Magstim 200. Results of stimulating in this fixed order have been shown to be similar to randomizing the intensities [14].

### Data analysis

For statistical analysis, we used repeated measures ANOVA. In addition to calculating the RMTs and input-output curves, the data was used to compute the latencies of MEPs with peak-to-peak amplitudes of 50 μV, 500 μV and 1mV, as done in previous studies [15]. The latency is defined as the time point where rectified EMG signals surpass a mean plus two standard deviations of the 100 ms pre-stimulus EMG level [16] [17]. The data were log-transformed [18] [19] [20] and the least-squares curve regression, which is a Gaussian-type curve with four parameters, was utilized to fit the data points of each participant individually [17] [21]. The slope of the IO curves was calculated from the tangent at the point where 50% of the maximal MEP size was reached. For two of the participants, who had a high threshold, we could not reach a plateau value for the IO curve, therefore these curves were excluded from the slope comparison.

## Results and discussion

The computational modeling, as well as the in-human results show that the PWM pulses approximate the neuronal effects of the conventional stimulus closely.

### Physiological response models

Figure 1(b) shows the change in membrane potential obtained from the RC model for both stimuli, with overall small dissimilarities. The modeled low-pass filtering properties of the neuron result in the membrane potential following the fundamental pulse frequency and attenuating the high frequency harmonics [22]. This dynamic of neural cells supports the principle of using PWM in TMS devices without causing unwanted side-effects due to the higher harmonics. Figure 1(c) displays the median excitation thresholds for both waveforms across the 2D cross-section of the pre-central crown, as obtained using the morphological neural models. The activation thresholds are consistently 5.6-6.2% lower for the pTMS2 pulse than for the Magstim pulse. The thresholds for each layer within the cortical hand muscle representation are shown in Figure 1(d), where each boxplot includes the data from five neuron clones within each layer. Linear regression between the thresholds for the two pulse types revealed a strong correlation (r^2^= 1.000, p= 0.000) with a slope of 0.939 (Figure 1(e)), indicating a consistently lower threshold for the pTMS2 pulses.

### MEP measurements

The RMTs as a percentage of the respective Magstim output are 41.34± 6.07% (mean ± standard deviation), and 38.00± 5.91% for pTMS2 stimuli, as shown in Figure 2(a). The pulse shape has a significant influence on the RMT (F_1.11_= 115, p< 0.01). Notably, the PWM pulses have a lower RMT than sinusoidal monophasic pulses for all participants (approximately 3%), as expected from the modeling results. The observed stronger effects of the PWM stimuli on the RMT may be related to the sharp edges and higher amplitude in the negative phase of the PWM pulses; other studies report similar results for rectangular pulses [15] [23]. However, further studies are required to confirm this.

**Figure 2.**
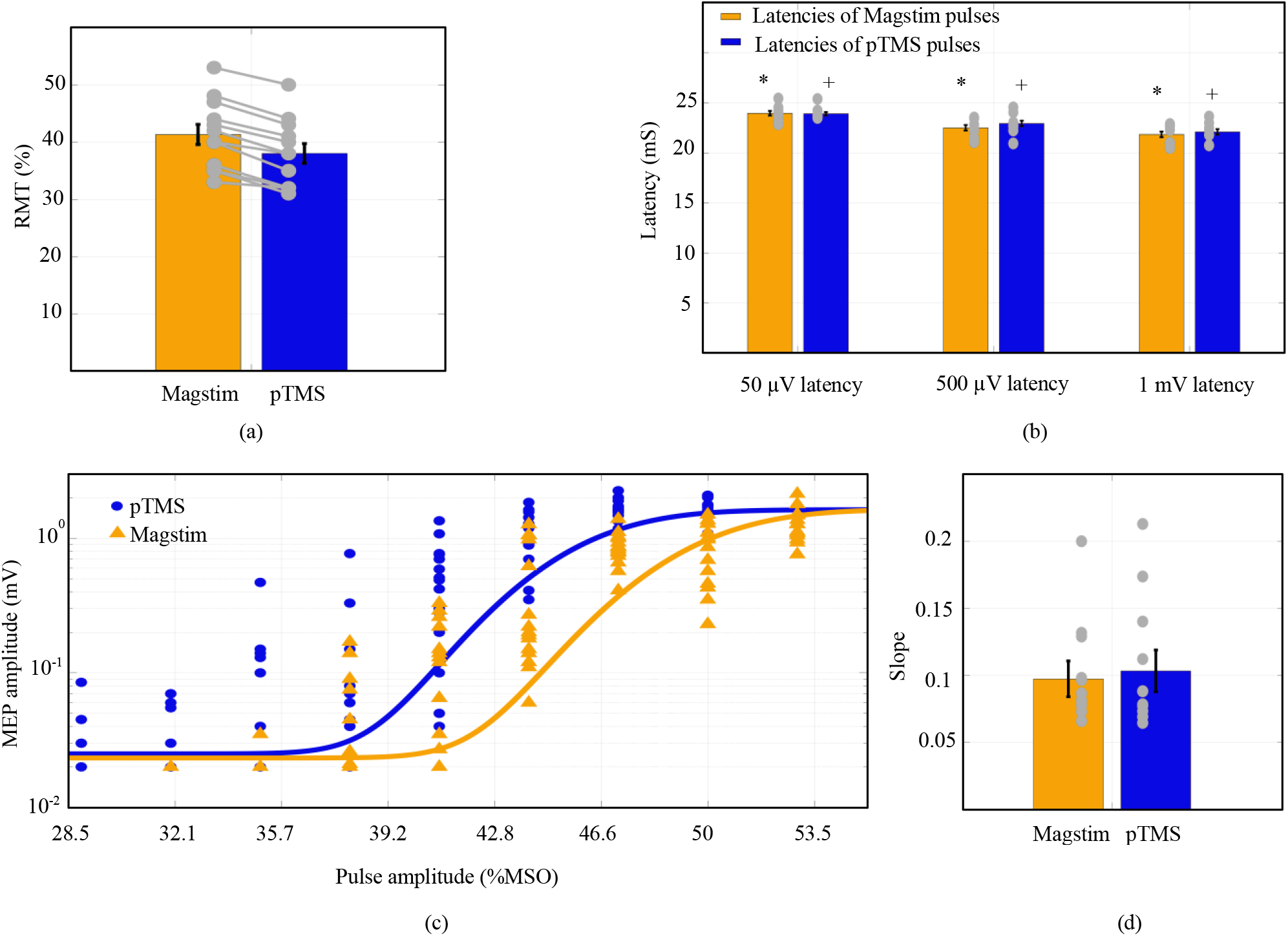
Results from twelve participants. (a) Measured resting motor thresholds of the conventional monophasic pulse and its modulation equivalent generated by the pTMS2 device. To calibrate the MSO of the two devices, the positive peak coil voltage of the pTMS2 device was compared with the Magstim device. (b) Average MEP latencies were measured for 50 μV, 500 μV and 1 mV peak-to-peak MEP amplitudes. * indicates the comparison between the MEP latencies across the different amplitudes for the Magstim pulses (*p* < .02), + indicates the comparison between the latencies across the different amplitudes for the pTMS2 pulses (*p* < 0.01), which is statistically significant for both devices. This significant difference indicates that different pulse sizes have different latencies. (c) Example IO curves of one participant for the Magstim pulse in orange and the pTMS pulse in blue, both in logarithmic scale (23-year-old female, RMT= 42% for Magstim and 38% for pTMS2 devices). (d) The IO curve slopes for both pulse types. For the IO curves, MEP measurements below 20 μV were set to 20 μV, as this was the lowest amplitude that was distinguishable from EMG signal noise. The MEP measurement was repeated 15 times for each amplitude, and the order of devices for the IO curves was counterbalanced to avoid order effects. For (a), (b) and (d), bars and whiskers show mean and standard error, respectively, with individual data points overlaid in grey.

The MEP latency is a reliable measure of the microcircuitry site of action potential initiation [15]. This latency is thought to show the number of synapses that the corticospinal volley crossed from the stimulation site to the target muscle. The MEP latencies for the two pulses are shown in Figure 2(b) which are not statistically significantly different between the devices (for 50 μV MEPs: F_1.11_= 0.07, p= 0.79, for 500 μV: F_1.11_= 0.65, p= 0.44, and for 1 mV: F_1.11_= 0.58, p= 0.46) while they differ for different MEP amplitudes. This supports the hypothesis that conventional and PWM pulses activate the same sites in the microcircuitry around the RMT value; Goetz et al. and D’Ostilio et al. have reported that different pulse shapes can cause different latencies and possibly activate different sites in the primary motor cortex or more than one population of axons [15] [17].

### IO curves

It has previously been reported that the stimulus shape affects the slope of the IO curve [17] [13]. Figure 2(c) shows an example of a sigmoidal IO curve of one participant for both devices. The raw EMG data for this participant is shown in Figure S5. Across the participants, the slopes of the IO curves are not significantly different between the devices (F_1.10_= 0.08, p= 0.77), as displayed in Figure 2(d). Together, the measured motor responses and IO curves indicate that the neural response to the conventional and PWM stimuli only differ by a small shift but not in their mechanisms of action.

### Side effects

No adverse events occurred during or after the stimulations. Participants did not report a subjective difference during stimulation, apart from a change in the sound emitted during pulse firing.

## Limitations

More research is needed to examine the brain’s response to different PWM pulse shapes, especially biphasic waveforms, and this study should be repeated for a larger participant cohort to replicate the findings. The pTMS2 device is currently limited to lower stimulation amplitudes than the Magstim 200^2^ for the pulse widths used here, which limited the data collection for individuals with very high thresholds. The maximum pulse amplitude of pTMS2 was 1600 V, compared to the maximum outputs for the Magstim Rapid, MagVenture MagPro, and Magstim 200^2^ which are approximately 1650, 1800 and 2800 V, respectively [15] [24]. Additionally, measurements of the clicking sound and electromagnetic noise are necessary for a better comparison of artifacts.

## Conclusion

TMS technologies are moving towards more programmable approaches to nerve stimulation. This study shows that PWM-based TMS can effectively imitate the effect of conventional stimuli on the cortex. Future applications of these TMS devices with new modulation paradigms might aid in finding new treatments for psychiatric and neurological diseases.

## Acknowledgements

This work was supported by a program grant from the MRC Brain Network Dynamics Unit at the University of Oxford (MRC MC_UU_0003/3), including an MRC iCASE fellowship, and supplemental funding to TD by the Royal Academy of Engineering. JO’S is supported by a Sir Henry Dale Fellowship from the Royal Society and the Wellcome Trust (HQR01720). This research was conducted in part in the Wellcome Centre for Integrative Neuroimaging, which is supported by core funding from the Wellcome Trust (203139/Z/16/Z). For the purpose of Open Access, the authors have applied a CC BY public copyright licence to any Author Accepted Manuscript version arising from this submission.

The authors would like to thank Magstim Company Ltd (UK) for providing the stimulation coil and valuable guidance on device design considerations.

## Conflict of interest and Financial disclosures

Prof. Denison reports personal fees from Medtronic and Bioinduction, outside the submitted work. Dr. Memarian and Ms Wendt report consulting fees from INBRAIN Neuroelectronics, outside the submitted work. For the pTMS device, a UK priority patent application (WO2021176219A1) was filed on 5^th^ March 2020.

## DATA AVAILABILITY STATEMENT

Data, and code used in this work are available in the public domain upon direct request from corresponding author.

## Authorship contributions

MMS and TD designed and built the pTMS2 device. MMS and KW conceptualized the study, performed the statistical analysis, and drafted the manuscript. KW conceptualized and performed the modeling analysis. MMS, KW, JO conceptualized and performed the in-human study. JO and TD provided funding for the study. All authors revised and approved the final version of the manuscript.

