## supplemental File for "Pulse-Width Modulation-based TMS mimics effects of conventional TMS on human primary motor cortex"

### SUPPLEMENTARY FILE

#### *Experimental measurements of pTMS2 device*

Representative measurement results of nine different stimulus waveforms are exhibited in Fig. S1. $V_{coil}$ and $I_{coil}$ were measured via a high-voltage differential probe (TA044, PICO TECHNOLOGY, UK) and a Rogowski current probe (I6000S FLEX-24, FLUKE, USA), respectively. The proposed two-cell architecture was tested with a cell link voltage of $V_{DC}$ = 800 V (peak-to-peak voltage 3.2 kV). For a 2.5 kHz cosine stimulus, the maximum energy delivered to the stimulation coil (D70 Remote coil, Magstim, UK) was measured to be 250 joules, which is equal to 100% of the maximum power of the Magstim rapid$^2$ option 2 equipment. As shown in Fig S1-S2, the proposed pTMS2 device can generate a double pulse and triple (called polyphasic) or a single pulse, in the form of biphasic and monophasic waves. The achievable pulse frequency starts at 2 kHz and can be increased up to 5 kHz.

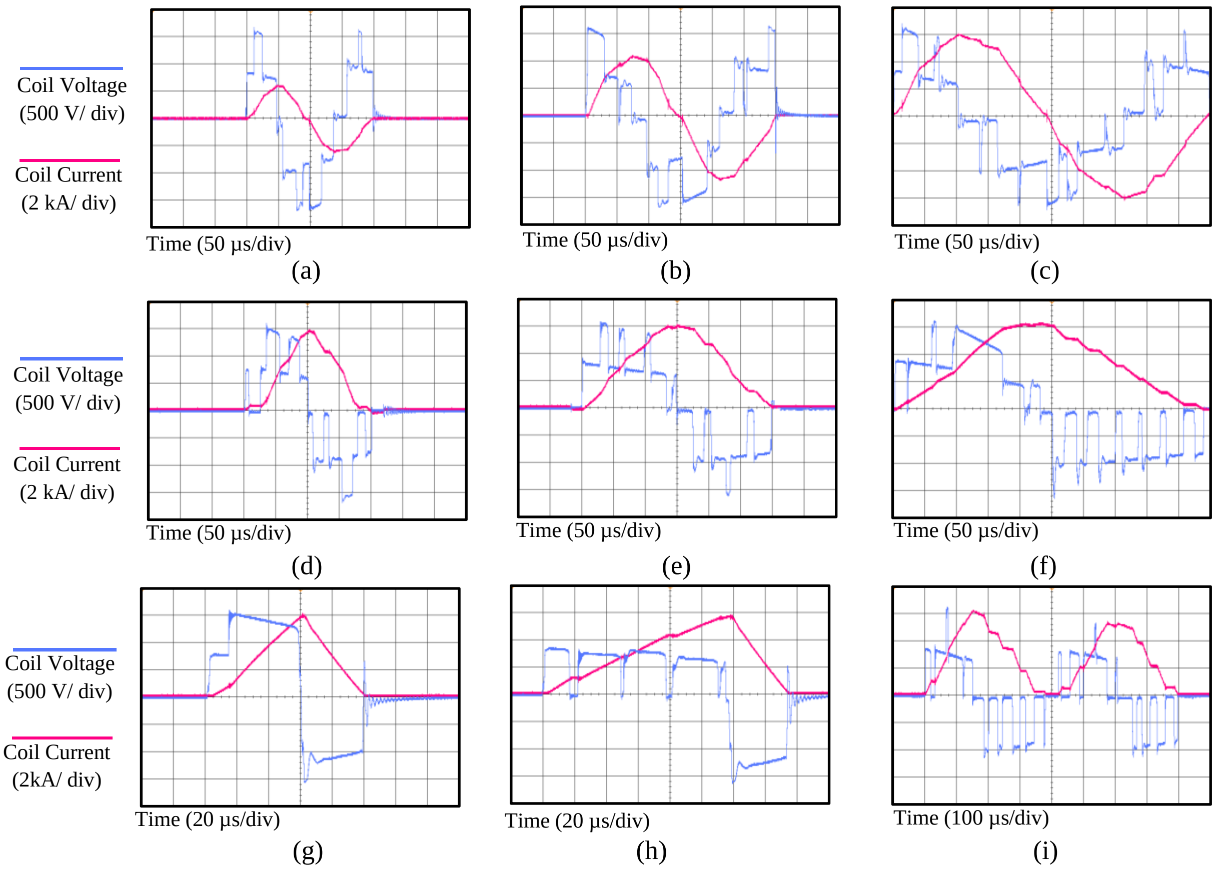


Figure S1 The measured waveforms for 9 different stimulus shapes: Voltage and current for (a) a 5 kHz cosine pulse, (b) a 3.33 kHz cosine pulse, (c) a 2 kHz cosine pulse, (d) a 5 kHz sine pulse, (e) a 3.33 kHz sine pulse, (f) a 2 kHz sine pulse, (g) a 60 μs rectangular stimulus (monophasic output, 60 μs positive and 20 μs negative phase, (h) a 120 μs rectangular stimulus (monophasic output, 120 μs positive and 40 μs negative phase) and (i) two continuous 2.5 kHz sine pulses. A digital oscilloscope with a sampling rate of 500 Ms/s was utilized in all measurements. No bandwidth restrictions or filters have been embedded to remove switching spikes.

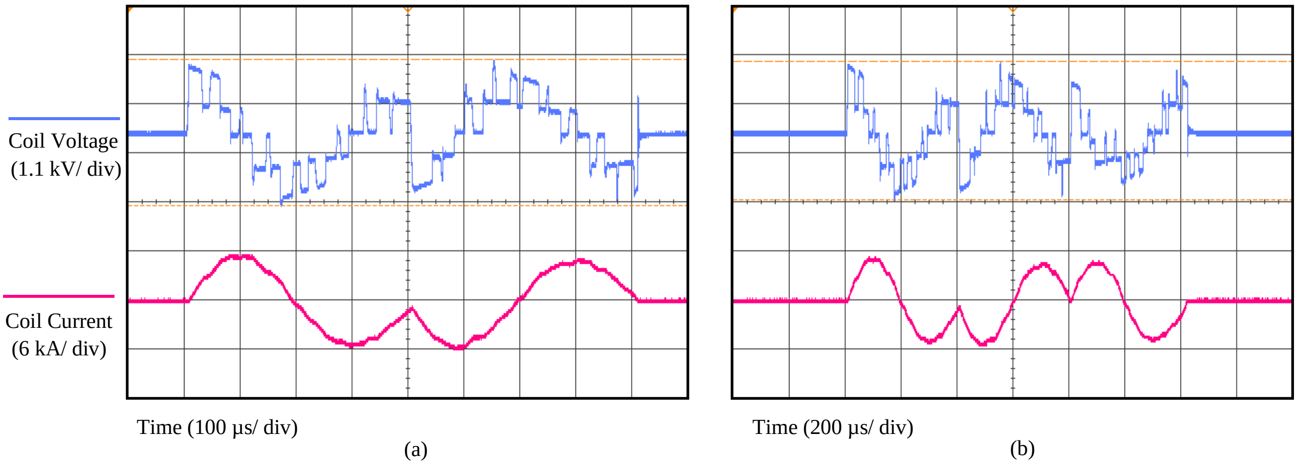


Figure S2 The measured waveform of the coil voltage and current for (a) double 2.5 kHz cosine stimuli where the two pulses have a phase difference of π radians and (b) triple 2.5 kHz cosine pulses with different phases.

*Efficiency of the pTMS device in generating repetitive pulses*

Repetitive TMS paradigms become gradually more popular modalities in basic neuroscience and for the treatment of several neurological and neuropsychiatric disorders. To enable further research on monophasic and biphasic rTMS methods, the pTMS device can generate high-frequency protocols with monophasic and biphasic or polyphasic stimulus shapes and repetition rates of up to 1 kHz (with an interstimulus interval of 1 ms). The energy recycling and the efficient modulation technique in the pTMS technology enable the generation of programmable and rapid stimulus sequences. In contrast to the previous TMS equipment, for which conventionally the output of several TMS devices must be combined, a single pTMS machine can generate and deliver a monophasic stimulus every 1 ms. Additionally, not all four waveforms must necessarily be the same, as it is possible to program and change the stimulus waveform, and the phase of every single pulse.

Fig. S3 illustrates examples of the proposed neurostimulator capability in generating rapid repetitive pulses. The measured parameters were set up for the generation of different sine and cosine stimuli with an interstimulus interval of 1 ms and different phases. Evidently, with a very short time interval, even lower than the required interval in the quadri-pulse stimulation (QPS) protocol, a consistent stimulus can be delivered. In addition, the regeneration mode of the inverter recovers significant energy from the coil to the capacitors [1]. The coil voltage and current measurement results for several classic repetitive protocols, such as 50 Hz and 100 Hz are provided in Fig. S4. The maximum stimulation intensity of each pulse in rTMS methods can be equal to the maximum system output (250 Joules). Therefore, in treatment protocols such as QPS, there is no need to use several devices and a combining module.

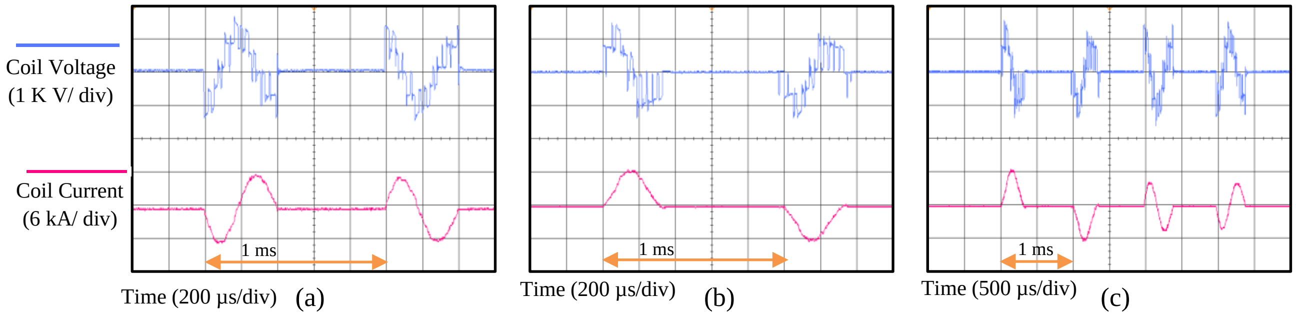


Figure S3 The measured waveforms for three different rapid rTMS protocol: Coil voltage and current for (a) two 2.5 kHz cosine pulses with a 1 ms interval, where the two pulses have a phase difference of π radians. (b) Two 2.5 kHz sine pulses with a 1 ms interval, where the two pulses have a phase difference of π radians. (c) Four 2.5 kHz sine and cosine pulses with different phases. The time intervals are 1 ms, which is 1.5 times lower than the minimum interval used in the QPS protocol. For the voltage graphs, each square represents 1 kV, and for the current graph, each square corresponds to 6 kA.

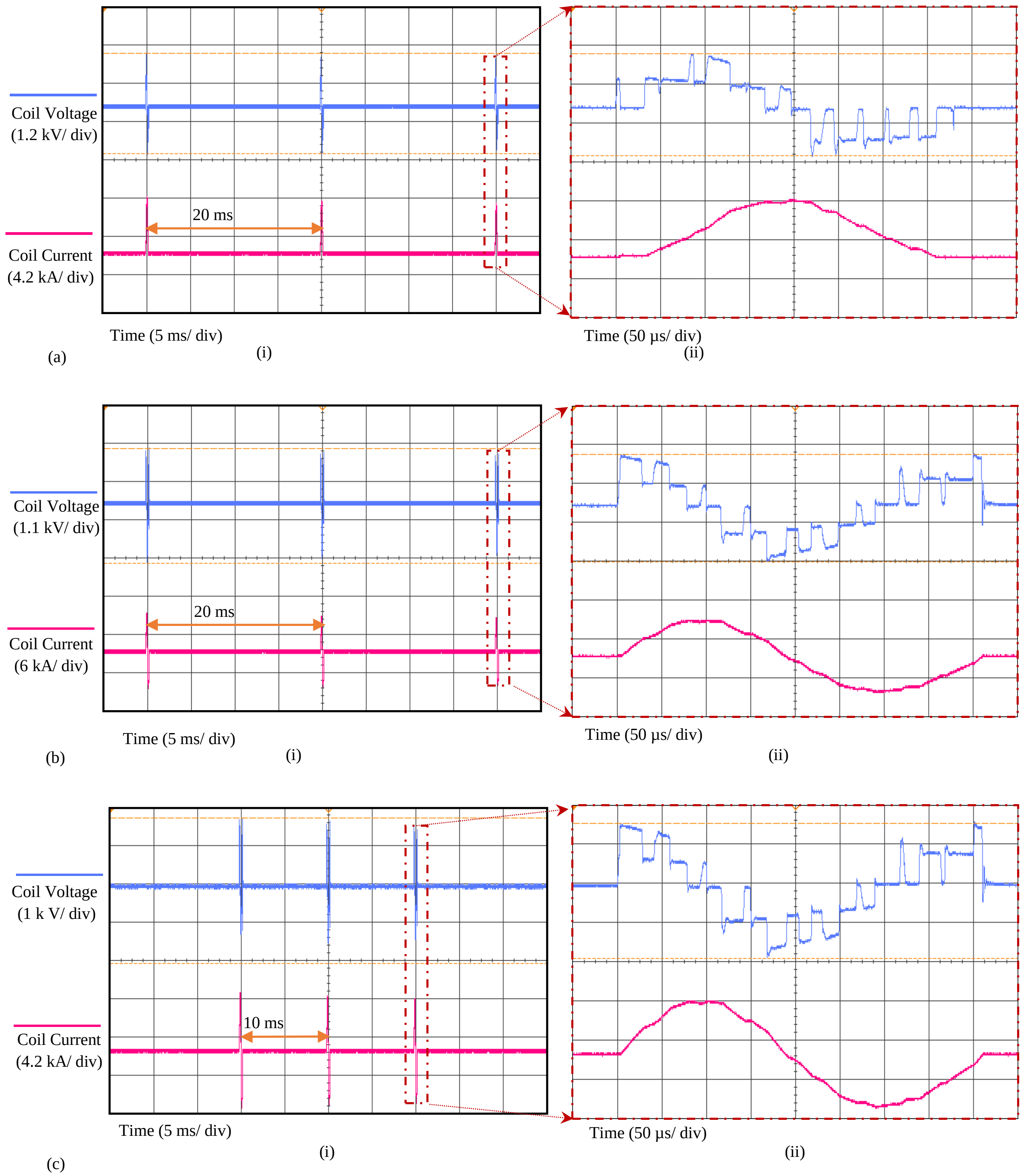


Figure S4 The measured waveform of the coil voltage and current for (a) the burst of 3 sine wave stimuli at 50 Hz (i.e., 20 ms between each stimulus), as a model of the theta burst stimulation (TBS) protocol with monophasic stimuli. (b) The burst of 3 cosine wave stimuli at 50 Hz, as a model of the classic TBS protocol with biphasic stimuli. (c) The burst of 3 biphasic stimuli at 100 Hz (i.e., 10 ms between each stimulus). In all figures, (i) indicates the overall view of the protocol and (ii) is zoomed in for one of the pulses of that protocol. For these measurements, the sampling rate of the digital oscilloscope is 10 Ms/s. No bandwidth restrictions or filters have been embedded to remove switching spikes.

*Morphological neural models*

The model relies on the assumption that the quasi-static approximation holds for neural stimulation, allowing the separation of the spatial and temporal components of the induced electric field. First, the spatial component of the electric field of a Magstim figure-8 coil was calculated using SimNIBS. Morphological models of neurons were then placed into the region of interest and the quasi-potentials at the model compartment centers calculated and applied to the neuron models as extracellular potentials. The temporal waveforms were simulated in Simulink using the stimulator circuits to replicate the stimulation pulses used in the in-human study. The extracellular potentials were then scaled by the temporal waveforms and used to calculate the membrane potential of each neuron compartment. A binary search algorithm was used to scale the coil current's rate of change at the pulse onset to find the activation thresholds of the neurons. Further details can be found in [2] [3]. The activation thresholds for PWM and conventional TMS pulses were compared.

*Recorded EMG values*

EMG recordings: The EMG signals were recorded using a Digitimer D440 Isolated Amplifier (Digitimer, Welwyn Garden City, UK), a Micro1401 (Cambridge Electronic Design, Cambridge, UK), a Digitimer HumBug Noise Eliminator and Signal version 7.01 (Cambridge Electronic Design, Cambridge, UK), with a 16-bit resolution at 10 kHz sampling rate, an amplifier gain of 1000 and a 10-1000 Hz filter (10-5000 Hz for one subject).

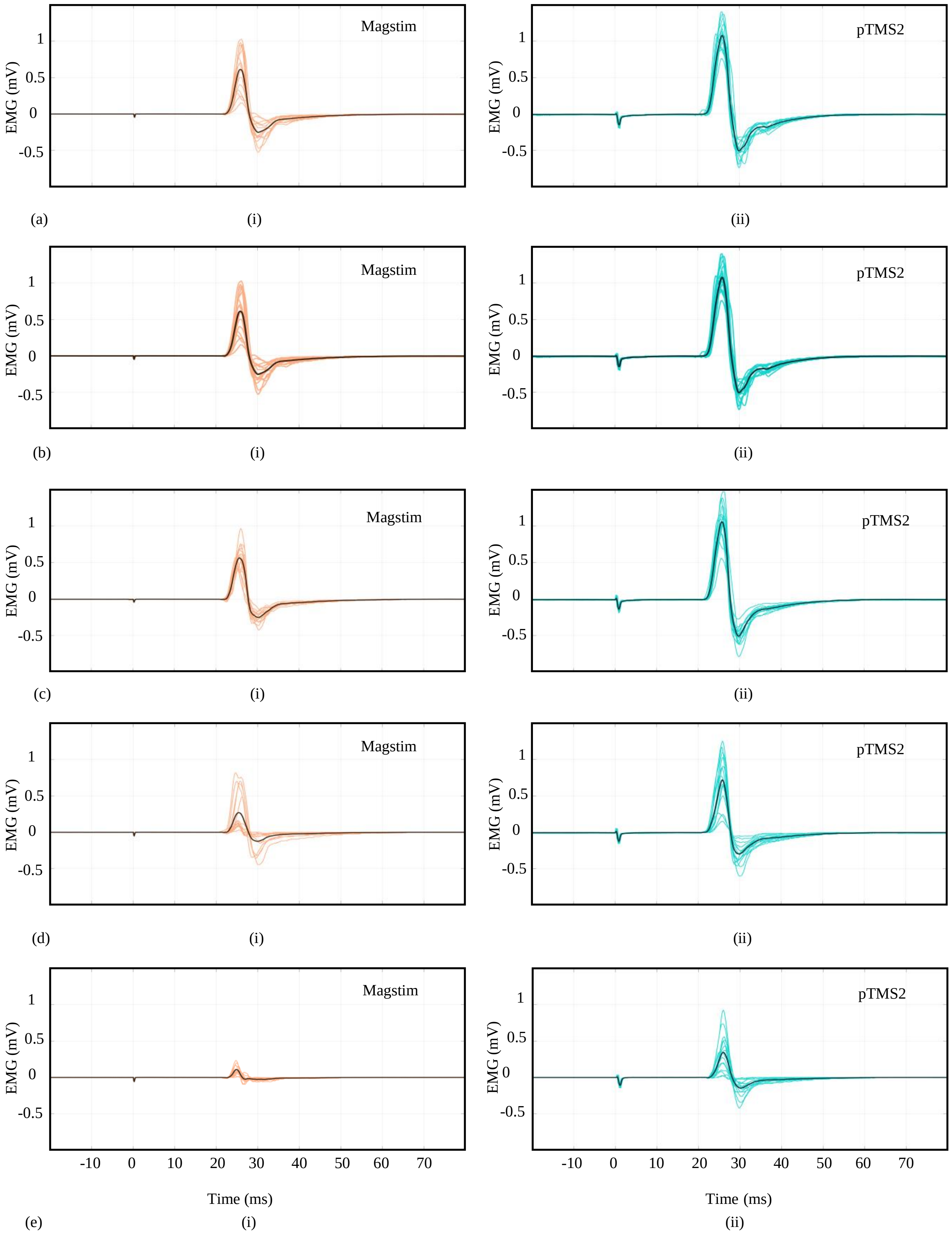
Magstim
pTMS2
EMG (mV)
1
0.5
0
-0.5
(a)
(i)
(ii)
(b)
(c)
(d)
(e)
-10
0
10
20
30
40
50
60
70
Time (ms)

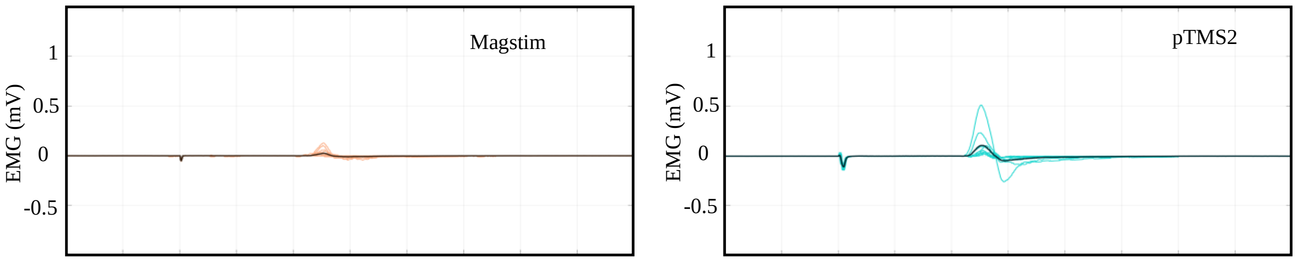


(f) (i) (ii)

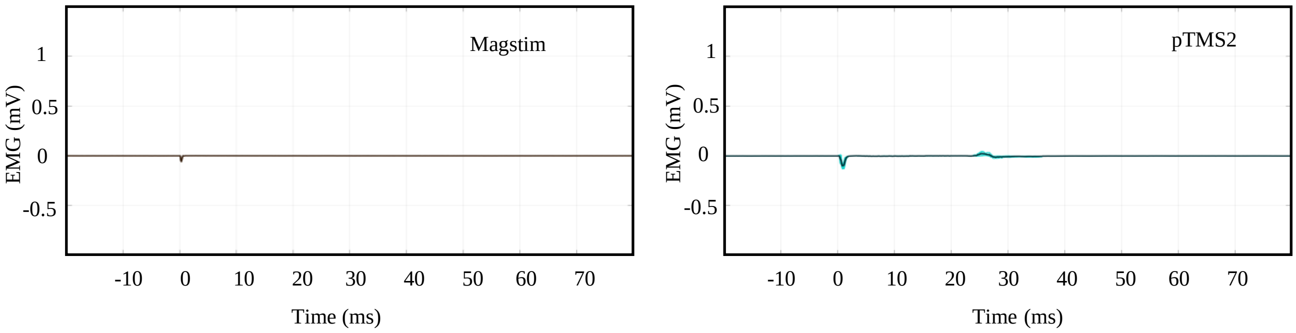


(g) (i) (ii)

Figure S5 Recorded EMG values for the subject whose IO curve is shown in Figure 1c. Stimulus intensity values are equal to (a) 50% of MSO, (b) 47% of MSO, (c) 44% of MSO, (d) 41% of MSO, (e) 38% of MSO, (f) 35% of MSO and (g) 32% of MSO of the Magstim 200. For each intensity, (i) shows the EMG response to the Magstim stimuli, (ii) shows the EMG response to the pTMS2 stimuli. The average EMG is shown by a black line.

#### *Individual input-output curves*

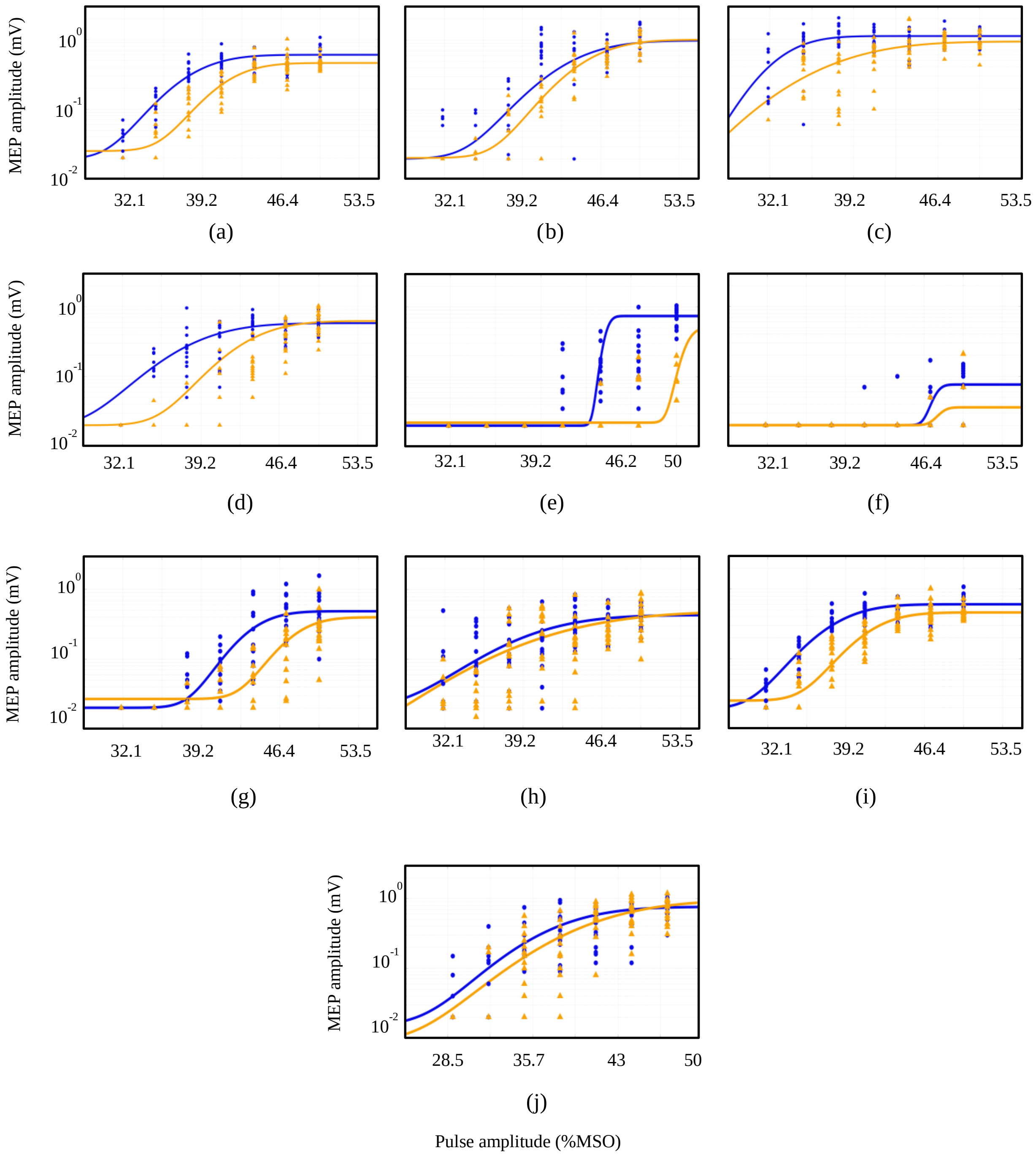


Figure S6 Individual input-output curves for the remaining participants for both devices. Each dot represents an individual data point, and the solid lines show the best line of fit. The vertical axis is on a logarithmic scale. The blue lines and dots are related to the pTMS device and the orange lines and dots are related to the Magstim device.
